# Variations in polarized trafficking of viral envelope proteins from insect-specific and insect-vectored viruses in insect midgut and salivary gland cells

**DOI:** 10.1101/2024.02.13.580057

**Authors:** Jeffrey J. Hodgson, Robin Y. Chen, Gary W. Blissard, Nicolas Buchon

## Abstract

Systemic viral infection of insects typically begins with primary infection of midgut epithelial cells (enterocytes) and subsequent transit of virus in an apical-to-basal orientation through the polarized enterocytes into the hemocoel. In the case of insect-vectored viruses, a similar yet oppositely oriented process (basal-to-apical virus transit) occurs upon secondary infection of salivary glands, and is necessary for virus transmission to non-insect hosts. To examine this inversely oriented virus transit in these polarized tissues, we assessed the intracellular trafficking of two model viral envelope proteins (baculovirus GP64 and vesicular stomatitis virus glycoprotein, VSV G) in the midgut and salivary gland cells of the model insect, *Drosophila melanogaster*. Using transgenic *Drosophila* fly lines that inducibly express either GP64 or VSV G, we found that both proteins were trafficked basally in midgut enterocytes. In salivary gland cells, VSV G was trafficked to apical membranes in most but not all cells, whereas GP64 was trafficked consistently to basal membranes. We further examined the mechanism of polarized trafficking in midgut and salivary gland epithelia and found that a cytoplasmic YxxØ motif in both VSV G and GP64 proteins is critical for basal trafficking of each envelope protein in midgut enterocytes, but dispensable for their trafficking in salivary gland epithelial cells. Using RNAi, we found that clathrin adapter protein complexes AP1 and AP3, as well as several Rab GTPases (Rab1, 4, 8, 10, 23, 30, and - 35), were involved in polarized VSV G trafficking in midgut enterocytes. Our results indicate that these viral envelope proteins encode the requisite information and require no other viral factors for appropriately polarized trafficking. In addition, they exploit tissue-specific differences in protein trafficking pathways to facilitate virus egress in the appropriate orientation for establishing systemic infections and vectoring infection to other hosts.

**Author Summary:** Viruses that use insects as hosts must navigate specific routes through the insect’s tissues to complete their life cycles. The routes may differ substantially depending on the life cycle of the virus. Some insect pathogenic viruses, such as baculoviruses, establish a systemic infection and this represents an endpoint in the infection cycle in the insect. In contrast, many insect-vectored viruses establish a systemic infection in the insect, but must also deliver infectious virus to the insect’s non-insect host. In both cases, the virus must first navigate through the midgut epithelium to establish a systemic infection, but insect-vectored viruses must also navigate through the salivary gland epithelium. Both midgut and salivary gland cells are polarized, and insect-vectored viruses appear to traffic in opposite directions in these two tissues. In this study, we asked whether two viral envelope proteins alone encode the signals necessary for polarized trafficking associated with their respective life cycles. Using two representative viral envelope proteins (VSV G and baculovirus GP64) and *Drosophila* as a model insect to examine tissue-specific polarized trafficking of viral envelope proteins, we identified one of the virus-encoded signals and several host proteins associated with regulating the polarized trafficking in the midgut epithelium.

## Introduction

To successfully establish a systemic infection in an animal host, most viruses must enter, replicate, and exit from various cell and tissue types. In some tissues, particularly those comprised of polarized cells, entry of the virus and egress of progeny virus particles must be coordinately directed to enable their successful infection of the organism. This is particularly important for arboviruses, viruses that are vectored between different host species. Of critical importance for arboviruses, the virus must transit through two insect tissue barriers: midgut and salivary gland epithelia.

Viruses that infect insects can be subdivided into a) those with simple life cycles requiring only one host to complete their life cycle (e.g., insect-specific viruses) and b) those with complex life cycles requiring more than one host (e.g., arthropod-borne viruses or arboviruses) and that utilize insects as vectors to infect vertebrate hosts. Some insect-specific viruses represent important components of ecosystems and may regulate insect populations in nature. One such group, baculoviruses, are virulent pathogens of insects and have been developed as commercial biopesticides to control important pest insect species [1–3]. Arthropod-transmitted diseases account for roughly 17% of infectious diseases worldwide, with over 3.9 billion people at risk for contracting arbovirus diseases [4–6]. Arboviruses are fascinating biologically because their life cycles are adapted to infect, replicate, and move between two highly divergent host groups, insects and vertebrates, posing substantial biological challenges.

In the case of both insect-specific viruses and arboviruses, insect infection is typically initiated when orally acquired virus particles infect midgut epithelial cells [7]. Viruses enter the polarized midgut cells (enterocytes) from the apical (gut lumen-facing) cell surface and establish a primary infection there. Following viral replication, progeny virions from the primary midgut infection are released from the basal surfaces of enterocytes into the open circulatory system (hemocoel) of the host insect, where virions disseminate and subsequently initiate a secondary round of infection that may include many other tissues. Thus, for systemic infections, the midgut epithelium is a critical early barrier, and efficient basal trafficking and escape are required to establish a robust secondary infection. In the case of arboviruses, secondary infection of the salivary glands is another pivotal component of their lifecycle that is required for transmitting the infection to vertebrates. In the salivary gland cells, the virus replicates and releases progeny virions into salivary secretions, which the insect vector injects into vertebrate hosts during blood feeding. Thus, appropriately navigating across the polarized cells of both the midgut and salivary glands is necessary to complete the arbovirus life cycle.

The midgut epithelium and salivary glands are structurally complex tissues [8–10] with dramatically different functions in the insect. Typical of most insects, the *Drosophila melanogaster* midgut is comprised of a single layer of polarized epithelial cells, containing four cell types: enterocytes (ECs), enteroendocrine cells (EEs), enteroblasts (EBs), and intestinal stem cells (ISCs) [11]. ECs represent the majority of the cells and biomass of the midgut, accounting for approximately 70% of the cells [12]. The midgut cells are protected apically from abrasion by a non-cellular chitinous structure called the peritrophic matrix (PM). The midgut cells are supported basally by a layer of extracellular matrix called the basal lamina and a layer of visceral muscles surrounding the gut. Midgut ECs typically have small apical finger-like projections (microvilli) on the apical surface facing the lumen of the gut. The basal surfaces of ECs comprise a tightly folded and relatively thin labyrinth of the basal membrane, which is closely apposed to the basal lamina and surrounded by visceral muscle [13]. Midgut cells secrete digestive enzymes into the gut lumen, and nutrients absorbed from the gut are transported across the midgut ECs and delivered to the hemocoel. The PM, midgut cells, and basal lamina also serve as a primary barrier against infection by microorganisms.

Similar to the midgut, the salivary glands are also comprised of a single monolayer of epithelial cells, and these cells produce salivary secretions. Salivary secretions facilitate digestion in insects in general and serve other specialized functions such as delivery of anticoagulants, anesthetics, and vasodilators in blood-feeding insects [14, 15]. Because salivary secretions may be required in large quantities on-demand during feeding, salivary secretions are often (e.g., in mosquitoes) stored in large cavities (acinar cavities) at the apex of each salivary gland cell. Release of the contents of these cavities into the salivary duct during feeding is carefully regulated [16]. Similar to the *Drosophila* adult midgut, which is divided into 5 morphologically and transcriptionally distinct regions, the adult salivary gland is divided into 3 regions that are hypothesized to be responsible for salivary secretion (medial and distal regions) and reabsorption (proximal region) [10, 16, 17]. The secretory cells of the medial salivary gland were analyzed in this study.

The mechanisms for directing polarized transport of proteins and budding of virus particles in invertebrate tissues (midgut, epidermis, Malpighian tubules, salivary glands, etc.) are largely unknown and may be similar or different in specific tissues. In one possible scenario, a virus may alter the host cell cytoskeleton and/or trafficking pathways to orient protein transport and viral release from polarized cells. In other cases, viral proteins may contain signals and motifs that interact with existing cellular infrastructure and traffic through intrinsic host pathways. Tissue-specific differences in these pathways (e.g., midgut enterocytes vs. salivary gland cells) may result in dramatically different protein and viral trafficking patterns. Directional budding of insect-specific or insect-vectored viruses has not been examined in detail in polarized insect tissues.

For many insect-specific viruses and arboviruses, virions enter midgut cells from the apical membranes adjacent to the midgut lumen, and progeny are released from basal membranes into the hemocoel, where they circulate and may infect a variety of other tissues. In the case of arboviruses, circulating virions then enter at the basal surfaces of salivary gland cells, replicate, and progeny virions are released from apical surfaces into apical invaginations that form cavities that interconnect with the salivary ducts. Virions in salivary secretions are then delivered to the vertebrate host upon blood feeding by the vector insect. This trafficking scheme represents an enigma: in the midgut, the virus must enter cells apically and release progeny virions basally, whereas in the salivary gland cells, the virus must enter basally and release progeny virions apically. Thus, it would appear that arboviruses either differentially modulate trafficking within midgut enterocytes vs. salivary gland cells or that cellular trafficking pathways may differ significantly in these specialized cell types. It is also plausible that some arboviruses may not rely on directional virion release from polarized cells and instead depend on sufficient levels of non-directional virion release. We hypothesize that the direction of polarized viral egress in a polarized cell is primarily determined by a combination of signals in the viral proteins and tissue-specific characteristics of the cell.

Polarized protein trafficking and virus budding involve many coordinated events, which may include: polarized trafficking of viral components for assembly of nucleocapsids at budding sites, polarized trafficking of pre-assembled nucleocapsids, polarized trafficking of the cellular fission machinery involved in deformation and fission of host membranes during budding, and polarized trafficking of viral envelope proteins. Because host membrane proteins can be dramatically segregated on apical or basal membranes of specific polarized cells, viral membrane proteins may utilize existing cellular differences in polarized trafficking pathways to facilitate trafficking to the appropriate polarized budding sites. Thus, there are many important questions regarding the roles of viral signals and host pathways that direct and regulate polarized protein trafficking and virion egress in these critical insect tissues.

In the current study, we used two model viral envelope proteins (baculovirus GP64 and vesicular stomatitis virus glycoprotein G, VSV G) as representatives of insect-specific viruses and arboviruses, respectively, to examine polarized trafficking in two vital insect tissues: midgut and salivary glands. Baculoviruses are virulent pathogens in many lepidopteran species, and their pathology primarily results from systemic infection of most host tissues [18]. VSV is an arbovirus that causes blister-like lesions in horses, cattle, swine, and occasionally in humans. VSV is transmitted to horses and cattle by blood-feeding insects such as biting midges, black flies, sand flies, and possibly mosquitoes [19]. For both viruses, the envelope proteins are essential for the production of infectious progeny virions [20, 21]. The VSV G protein has been used as a model for protein trafficking in mammalian cells [22], and VSV G is known to be specifically targeted to basal membranes of polarized mammalian MDCK cells [23, 24]. However, little or nothing is known regarding the trafficking of VSV G in the insect midgut. The structure and function of the baculovirus GP64 protein have also been studied extensively in the context of cultured cell infections [20, 25, 26], and basolateral GP64 localization was previously reported in the insect midgut in the context of baculovirus AcMNPV-infected enterocytes [27]. Thus, VSV G and GP64 represent highly relevant models for studies of viral envelope protein trafficking in polarized insect midgut and salivary gland cells.

To investigate viral envelope protein trafficking in these polarized tissues, we used the tractable model insect, *Drosophila melanogaster*, to generate a transgenic system for expressing and monitoring viral envelope protein trafficking. We found that expression of either GP64 or VSV G alone was sufficient for the trafficking of either protein to the basal membranes of midgut enterocytes. In salivary gland cells, GP64 was trafficked to basal membranes. However, in contrast to GP64, VSV G was trafficked to apical membranes in most salivary gland cells. To explore the mechanisms responsible for polarized trafficking in these key tissues, we first examined a previously identified YxxØ basal trafficking motif that is present in both VSV G and GP64. The conservation of the YxxØ motif in insect-specific and insect-vectored viral envelope proteins suggests that it could play an important role in the lifecycles of these viruses. Because the YxxØ motif interacts with clathrin adapter protein (AP) complexes, and these complexes are highly conserved from yeast to vertebrates, we focused on VSV G trafficking and used RNAi knockdowns to examine the role of clathrin adapter complexes in polarized trafficking in midgut enterocytes. In addition, because Rab GTPases are known regulators of vesicular protein trafficking, we also analyzed similarities and differences in the localization of 30 *Drosophila* Rab GTPases in these tissues. We then used RNAi to examine selected Rab GTPases for their potential roles in modulating or regulating polarized trafficking of VSV G in the midgut epithelium. We found that both VSV G and GP64 were trafficked basally when each was expressed alone in transgenic fly enterocytes, i.e., in the absence of infection or any other viral proteins. Alanine substitutions in the YxxØ motif disrupted basal trafficking of both VSV G and GP64 in midgut enterocytes. However, ablation of the YxxØ motif had no apparent effect on the trafficking of either VSV G or GP64 in salivary gland cells. Using RNAi knockdowns, we also found that AP-1 and AP-3 clathrin adaptor complexes and several Rab GTPases were important for basal VSV G trafficking in enterocytes.

## Materials and Methods

### Fly transgenesis

*Drosophila* fly lines encoding either WT VSV G (VSV G^WT^), WT GP64 (GP64^WT^), modified VSV G (VSV G^ΔY^) or modified GP64 (GP64^ΔY^) were generated by inserting the wt or modified ORFs downstream of the upstream activation sequence (UAS) in the *Drosophila* transformation plasmid pUASt (Drosophila Genomics Resource Center plasmid #1000). The VSV G^WT^ open-reading frame (ORF) was PCR amplified from VSVG-BP95NOTSV (kindly provided by F. Boyce) using primers: forward 5’AAA*GAATTC*TGACACTATGAAGTGCCTTTTGTACTTAGC3’ and reverse: 5’AAA*TCTAGA*TTACTTTCCAAGTCGGTTCATCTC3’ and cloned into EcoRI/XbaI sites of pUASt, generating pUAST-VSV G. The modified VSV G^ΔY^ ORF contained Y501a and I504a substitutions and was generated by PCR from the VSV G^WT^ template, using the Forward primer (containing the EcoRI site, in italics): 5’aaa*<u>gaattc</u>*TGACACTATGAAGTGCCTTTTGTACTTAGC3’ and the reverse primer: 5’CGGTTCATCTC**agc**GTCTGT**agc**AATCTGTCTTTTCTTGGTGTGC3’ (mutagenic codons in lower case bold). This primary PCR amplicon was used directly as template for a secondary PCR using the same forward primer: 5’aaa*gaattc*TGACACTATGAAGTGCCTTTTGTACTTAGC3’ and a reverse primer containing an XbaI site (italics): 5’aaa*<u>tctaga</u>*TTACTTTCCAAGTCGGTTCATCTC**agc**GTCTG3’, then EcoRI/XbaI cloned into pUASt to generate pUASt-VSV G^ΔY^. The GP64^WT^ ORF was PCR amplified from AcMNPV genomic DNA using the forward primer (containing an XbaI site, in italics): 5’aaa*tctaga*ATGGTAAGCGCTATTGTTTTATATGTGC3’ and the reverse primer (containing an XbaI site, in italics): 5’aaa*tctaga*ATATTGTCTATTACGGTTTCTAATCATACAG3’ and XbaI-cloned into pUASt to generate pUASt-GP64. The mutant GP64^ΔY^ ORF, encoding the Y502a and I505a codon substitutions, was generated by PCR. A primary PCR amplicon was generated from the pUASt-GP64^WT^ template using the forward primer (containing an XbaI site, in italics): 5’aaa*<u>tctaga</u>*ATGGTAAGCGCTATTGTTTTATATGTGC3’ and the reverse primer: 5’**agc**CATACA**agc**CAAAAATAAAATCACAATTAATATAATTACAAAGTTAACTAC3’ that introduced both alanine codons (bold). This primary PCR amplicon was used directly as template for a secondary PCR using the same forward primer: 5’aaa*tctaga*ATGGTAAGCGCTATTGTTTTATATGTGC3’ and the reverse primer (containing an XbaI site, in italics): 5’aaa*<u>tctaga</u>*TTAATATTGTCTATTACGGTTTCT**agc**CATACA**agc**CAAAAATAAAATCAC3’ (substituted codons in bold). High fidelity KOD DNA polymerase (Takara) was used to amplify the entire VSV G and GP64 ORFs (WT and ΔY) which were verified by Sanger sequencing. The resulting pUASt-based plasmids containing the VSV G or GP64 fragments were injected into W1118 flies (BestGene Inc, California), and transgenic fly lines were verified for expression of the viral envelope proteins (WT and ΔY versions of VSV G and GP64) by immunoblot analysis. A single transgenic line (expressing each WT or ΔY version of VSV G or GP64) was selected and used for all experiments. The 3’ ends of the VSV G and GP64 ORFs were PCR-amplified (using Taq polymerase, NEB) from DNA isolated from each fly line, and the PCR amplicons were sequenced to verify the WT and ΔY forms of cytoplasmic domain sequences for each line from VSV G or GP64.

### Fly husbandry and genotypes

Localization patterns of the WT and alanine-substituted forms of VSV G and GP64 or YFP-Rab GTPases in *Drosophila* midgut enterocytes and salivary gland cells were assessed in the F1 offspring of *hs-hid; da-Gal4, tub-Gal80^ts^* female ubiquitous driver flies crossed with male flies carrying WT or modified *UAS-GP64 or UAS-VSV G*. The same driver flies were crossed with the various UAS-regulated WT and alanine-substituted viral envelope proteins or YFP-Rab transgenes. For assessing Rab GTPase localization patterns or to determine total levels of WT and alanine-substituted VSV G in permeabilized and non-permeabilized tissues, we used F1 progeny from *hs-hid*; *da-Gal4; +* ubiquitous female driver fly matings. *Myo-Gal4, UAS-nlsGFP, tub-Gal80^ts^; UAS-VSV G^WT^,* and *Myo-Gal4, UAS-nlsGFP, tub-Gal80^ts^; UAS-GP64^WT^* midgut-specific female driver lines were crossed with male RNAi and control lines for quantifying VSV G basal-to-apical distribution in the midgut enterocytes. The F1 progeny were allowed to develop at 18°C under a 12L:12D light cycle on an artificial diet (Table S1). F1 adults (3 to 7 days post-emergence) were then incubated at 29° C for 3 or 5 days to induce transgene expression prior to tissue dissection for processing and microscopy. The incubation time at 29°C in each case was carefully selected to avoid gut dysplasia while maximizing the duration of RNAi induction [28]. Control AttP2 (36303), AttP40 (36304), and UAS-regulated RNAi lines were obtained from the Bloomington *Drosophila* Stock Center (BDSC): Rab1 (34670). Rab4 (33757), Rab8 (34373), Rab10 (26289), Rab11 (27730), Rab23 (55352), Rab30 (31120), Rab35 (80547), AP-1µ (27534), AP-1,2β (28328), AP-2μ (28040), and AP-3μ/cm (27282). A fly line ubiquitously expressing E-Cadherin-GFP used in the cell polarity markers analysis was generously provided by Bruce Edgar (University of Utah).

### Immunostaining and Microscopy

Flies were anesthetized on ice before brief immersion in 70% ethanol to remove cuticular hydrocarbons, then transferred to PBS (pH 7.4) in a 9-well glass spot plate. Midguts and salivary glands were dissected in PBS and immediately transferred to 1 ml of 4% paraformaldehyde in PBS (pH 7.4) at RT for 1 h of fixation. Tissues were rinsed (3x, 10 min each at RT) with 1 ml of PBS (for non-permeabilized tissues) or PBS-T (PBS pH 7.4 containing 0.1% Triton X-100) for cell permeabilization and blocked in 3% BSA in PBS (pH 7.4) for a minimum of 3 h before incubation with primary antibodies (1:1000) in 1% BSA in PBS overnight at RT. Primary antibodies were removed and tissues were washed (3x, 10 min each at RT) with 1 ml PBS. Washed tissues were incubated in secondary antibodies (1:1000) and Alexa Fluor 555 phalloidin (1:1000) (Invitrogen) in 1% BSA in PBS (pH 7.4) for 2-4 h in darkness at RT. Secondary antibodies were removed, and tissues were washed (3x, 10 min each at RT) with PBS, then stained in 1 ml of 0.5 µg/ml DAPI (Sigma-Aldrich) in PBS for 30 min at RT in darkness, then washed (3x for 10 min at RT) with PBS. Stained tissues were mounted onto slides in glycerol-based, aqueous mounting media containing an antifadent (Citifluor AF3). Tissues in mounting media were placed between pedestals consisting of two layers of double-sided Scotch tape to prevent crushing of the tissues by the coverslip. The enterocytes of midgut region 2 and the secretory cells of the medial salivary gland were analyzed in this study. Slides were imaged on a Zeiss 710 confocal microscope using the 63x oil immersion objective. Primary antibodies used for immunostaining were directed against: AcMNPV GP64 (AcV5), VSV G (8G5F11, gift from Gary Whittaker, Cornell University), Discs large (DSHB 4F3), GFP (Invitrogen, A10262), Integrin β1 (DSHB 7E2), Snakeskin (Gift from Mikio Furuse, NIPS), and Tetraspanin 2A (Gift from Mikio Furuse, NIPS). Alexa Fluor 647 donkey anti-mouse, Alexa Fluor 555 donkey anti-rabbit, and Alexa Fluor 488 goat anti-chicken secondary antibodies (Invitrogen) were used in this study.

### VSV G and GP64 protein distribution in fly tissues

VSV G and GP64 signal distributions in immunostained tissues were determined using ImageJ (v.1.53c). To determine the relative basal-to-apical distributions of WT and modified VSV G and GP64 constructs, individual cells were assessed for the mean gray values of protein staining in a rectangular region of interest (ROI) adjusted to the width of each cell along the basal-apical axis of the cell. The polarized orientations of midgut or salivary gland cells were identified by the positions of phalloidin-stained actin in the basal visceral muscles and the apical brush border, and by differential interference contrast (DIC) microscopy. In some instances, when midgut cells were slightly curved, a segmented (curved) ROI was selected to account for the cell curvature. The same ROI was used to measure the mean gray values of the immunostained VSV G or GP64 (Alexa 647) and nuclear (DAPI) signals. Because cells were not uniform in size or staining intensity, the basal-to-apical distance was expressed in percentage values, and the signal (VSV G, GP64, and DAPI) intensity within each cell was standardized based on the average signal intensity within that cell. Each cell’s resulting standardized gray values were binned in 5% intervals across the basal-apical axis (to account for differences in cell sizes), averaged across all cells for each condition, and plotted onto graphs. The percentage of total viral protein signal in the apical (top 40%) and basolateral (bottom 20%) compartments of enterocytes and salivary gland cells were quantified from these data to determine the effects of the tyrosine motif mutation and various RNAi of host trafficking components on viral protein distribution in these compartments. These compartments in enterocytes were selected based on DLG and DAPI staining to represent areas of the apical and basolateral compartments free from any interference of the nucleus where the viral protein is absent (See schematic in Fig. S1). Distinguishing between the apical and basolateral compartments in *Drosophila* salivary gland cells is difficult due to their unique structure [i.e., extensive apical membrane invaginations (canaliculi) reaching deep inside the cell (Fig. 1)], and therefore could not be accurately determined with our current method. Since these invaginations rarely extend past the nuclei, as indicated by phalloidin staining of the actin-rich canaliculi, we decided to quantify viral protein signal in the bottom (basal) 20% of the salivary gland cells to minimize any influence of the nucleus or canaliculi (see Fig.1C, 1D, and S1). The master gain of the laser for acquiring VSV G or GP64 and nuclear stain signals was optimized for each image to avoid pixel saturation. Data shown was compiled from 3 independent experiments, in which ≥20 cells from each of at least 3 individual midguts or salivary glands (total N ≥ 69) were assessed per experiment. The total number of cells used for each quantification is noted in the figures.

**Figure 1.**
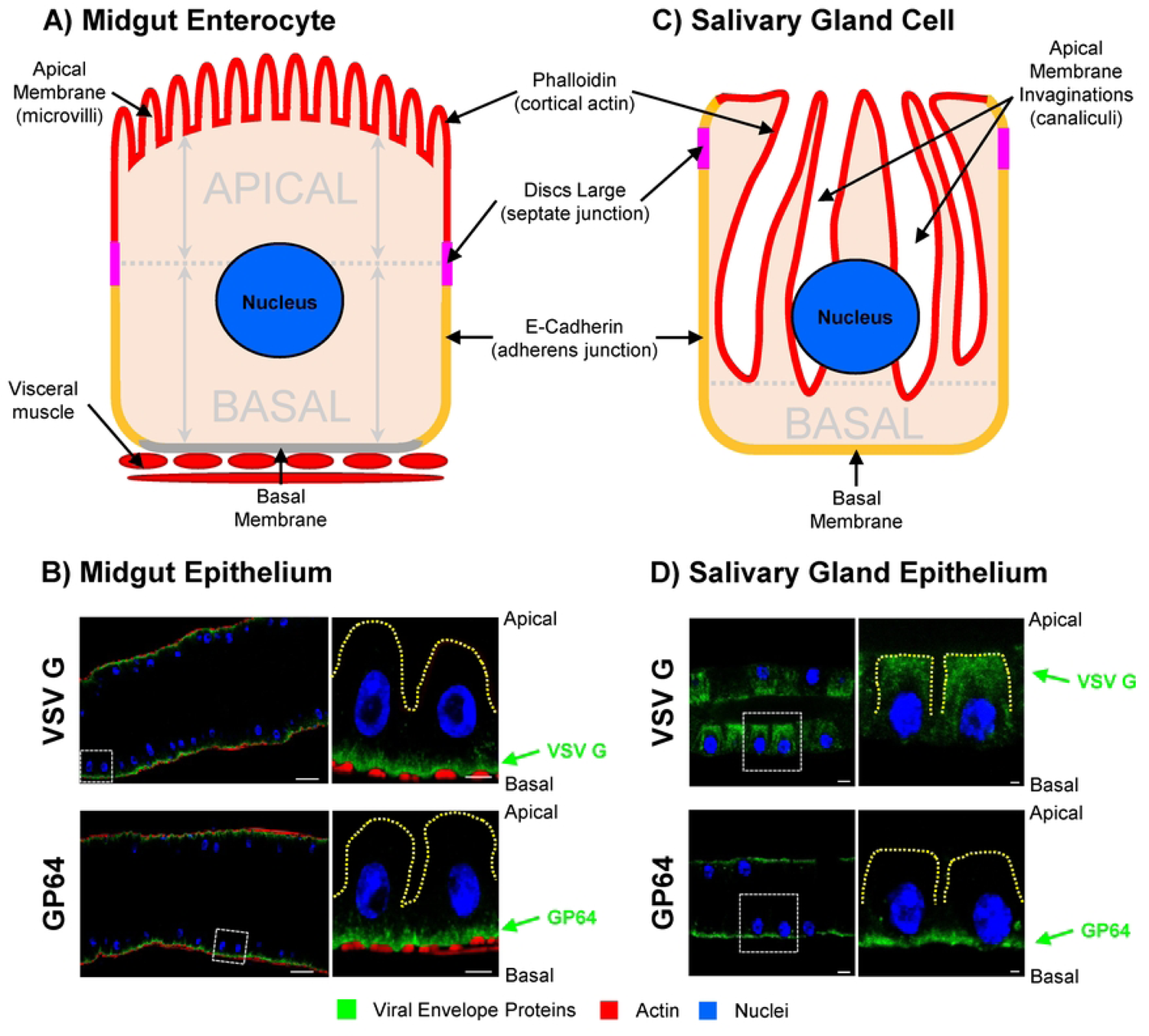
Localizations of VSV G and GP64 in *Drosophila* midgut and salivary gland in the absence of VSV or baculovirus infection. (A) Schematic representation of a *Drosophila* midgut enterocyte showing polarized morphology with apical microvilli (red) adjacent to the gut lumen and basal membrane (gray) adjacent to the visceral muscles. The relative locations of the nuclei (blue) and polarity marker proteins for septate junctions (Discs large; magenta) and adherens junctions (E-Cadherin; orange) are indicated. B) Localization of VSV G (top panels) and GP64 (bottom panels) ectopically expressed in adult *Drosophila* midgut (left) and associated enterocytes (dashed box, right). Both viral envelope proteins (green) are enriched basally adjacent to visceral muscles (red). C) Schematic representation of an adult *Drosophila* salivary gland epithelial cell showing deep invaginations of the apical membrane (canaliculi; red). The relative locations of the nuclei (blue) and polarity marker proteins for septate junctions (Discs large; magenta) and adherens junctions (E-Cadherin; orange) are indicated. D) Localization of VSV G (top panels) and GP64 (bottom panels) in adult *Drosophila* salivary gland (left) and associated epithelial cells (dashed box, right). VSV G was enriched apically or distributed throughout the cell, whereas GP64 was consistently basal. Tissues were dissected from adult *Drosophila* expressing each viral envelope protein ubiquitously under the *da-Gal4 tub-Gal80^ts^* driver. At 3 to 7 days post-eclosion, flies developed at 18° C were shifted to 29° C for 3 days to induce expression of VSV G or GP64. Viral envelope proteins (green) were labelled with mouse anti-VSV G or anti-GP64 primary antibodies and Alexa Fluor 647 donkey anti-mouse secondary antibody, actin (red) was labelled with Alexa Fluor 555 phalloidin, and nuclei (blue) were stained with DAPI. Scale bars are 25 µm for tissue images (left) and 5 µm (midgut) or 10 µm (salivary gland) for cellular images (right).

To assess the total levels of VSV G^WT^ and VSV G^ΔY^ staining in cells, the Integrated Density values (IntDen = Mean Gray × Area) were determined in ROIs drawn around entire cells (guided by actin staining and/or DIC images) using the freehand tool in ImageJ. To measure the basal surface levels of VSV G in non-permeabilized tissues, a standardized ROI (approximately the width of a cell) was used for individual basal cell membrane measurements. The master gain of the laser for acquiring VSV G stain signals was kept constant (while avoiding pixel saturation) for groups of WT, ΔY, or negative (N) samples of both the permeabilized and non-permeabilized tissues. The numbers of cells used for each quantification of total VSV G levels in enterocytes were: VSV G^WT^ permeabilized (n = 44), VSV G^ΔY^ permeabilized (n = 40), VSV G^WT^ non-permeabilized (n = 40), and VSV G^ΔY^ non-permeabilized (n = 35).

### Statistical analysis

All statistical analyses were conducted using R Statistical Software (RStudio v.4.1.2) [29]. Data for each condition was subjected to the Shapiro–Wilk test to determine whether it was normally distributed. For single comparisons of conditions, Welch’s t-test was used to compare normally distributed data, while the Wilcoxon rank-sum test was used to compare data that were not normally distributed. For multiple comparisons of conditions, one-way analysis of variance (ANOVA) was used to compare normally distributed data, while the Kruskal–Wallis test was used to compare data that were not normally distributed. If statistically significant differences overall were found (p ≤ 0.05), then a *post hoc* multiple comparison analysis (with Bonferroni p-value correction) was performed to identify which specific conditions were significantly different from each other.

## Results

### GP64 and VSV G are trafficked to basal membranes in *Drosophila* midgut enterocytes independent of viral infection

For viruses that are acquired orally by insects, the midgut is a cellular barrier to systemic infection. Following replication in the midgut epithelial cells (i.e. enterocytes or ECs), egress of progeny virions from this primary site of infection into the hemocoel is critical for initiating the systemic secondary phase of infection. Envelope proteins are crucial viral structural components since they are often required for virion assembly and egress, and are necessary for virion binding and entry into host cells [30, 31]. Therefore, for many insect viruses, transport of viral envelope proteins to basal membranes of infected midgut enterocytes is essential for efficient assembly and egress of infectious progeny virions. However, it remains unknown whether this requires viral reprogramming of the trafficking pathways in enterocytes, whether these viral proteins utilize existing cell trafficking machinery, or whether they do indeed accumulate basally. We selected two model viral envelope proteins to examine this phenomenon in the insect midgut: GP64 as a representative envelope protein from insect-specific viruses, and VSV G as a representative envelope protein from viruses vectored by insects (arboviruses) [32, 33]. GP64 is one of the more intensively studied envelope proteins from insect-specific viruses, and VSV G has served as an important model protein in numerous membrane protein trafficking studies in mammalian cells [34–36].

We first asked whether GP64 or VSV G would traffic basally in insect midgut enterocytes, in the absence of infection. To address this question, we generated transgenic *Drosophila* lines that inducibly express either VSV G or GP64, without any other viral protein(s) that might modify cellular architecture or pathways. We found that both VSV G and GP64 were concentrated in the basal portion of the cells adjacent to the visceral muscles (Fig. 1A, B). The basal-most 20% region in the enterocytes accounted for 70.4 ± 1.28% (mean ± SE, n = 96) and 70.3 ± 1.25% (n = 90) of the total VSV G and GP64 signal, respectively (Fig. 2B, VSV G and GP64, WT). In addition, we confirmed surface display of VSV G on basal membranes of enterocytes by immunostaining non-permeabilized tissue (Fig. S2). Therefore, the information necessary for basal trafficking in insect midgut enterocytes is encoded in both viral envelope proteins and does not require other viral proteins or infection.

**Figure 2.**
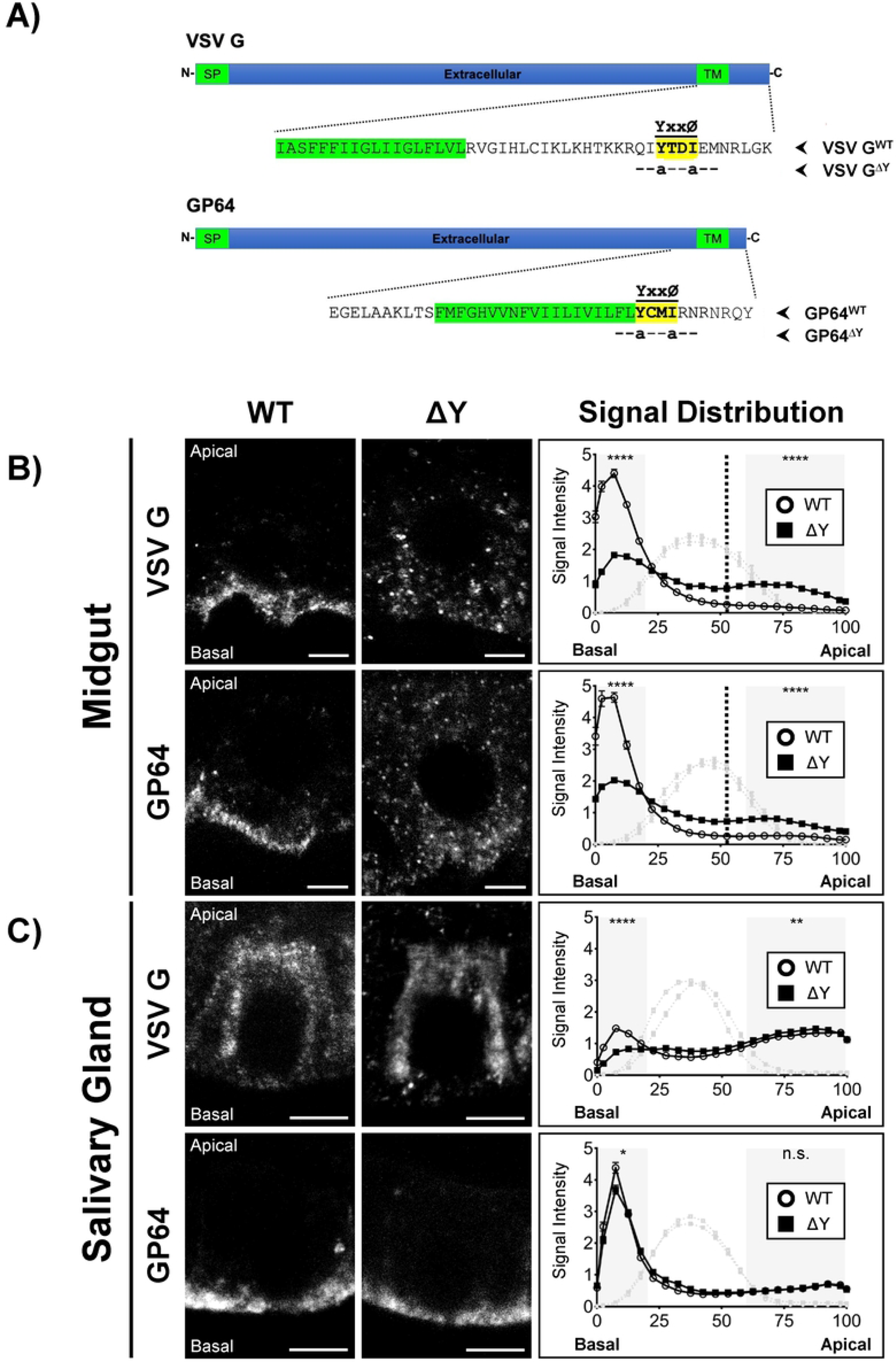
Analysis of the role(s) of VSV G and GP64 cytoplasmic YxxØ motifs for envelope protein trafficking in *Drosophila* midgut enterocytes and salivary gland cells. A) Schematic illustration of the cytoplasmic tail of VSV G and GP64, showing the location of the YxxØ motif (yellow highlight) in each protein and the engineered alanine substitutions (lower case “a”). B) Representative images showing the distribution of WT (left) and alanine-substituted (right) viral envelope proteins (VSV G and GP64) in midgut enterocytes after 3 days of ubiquitous expression under the *da-Gal4 tub-Gal80^ts^* driver. A graph comparing the average relative distributions (basal-to-apical) of each protein (WT vs. ΔY) is shown on the right of each pair of microscopic images. While VSV G^WT^ and GP64^WT^ showed strong basal enrichment (open circles), VSV G^ΔY^ and GP64^ΔY^ were more evenly distributed in midgut enterocytes (closed squares). The average relative position of the nucleus (determined by DAPI signal intensity) was also plotted as a light gray line in the background of each graph to indicate areas relatively free from its influence (shaded gray areas). The basal and apical regions on the graphs were established based on the location of the septate junction (discs-large polarity marker) (Fig. S1), and the division is indicated at 52.5% on the x-axis by a vertical dashed line. (C) Representative images show the distribution of WT (left) and modified (right, ΔY) viral envelope proteins (VSV G and GP64) in salivary gland cells. As above, a graph comparing the average relative distributions (basal-to-apical) of each protein (WT vs. ΔY) is shown on the right of each pair of microscopy images. Both VSV G^WT^ and VSV G^ΔY^ were distributed near or adjacent to the canaliculi in salivary gland cells. Both GP64^WT^ and GP64^ΔY^ were localized to the basal region in salivary gland cells. The average relative position of the nucleus was also plotted as a gray line. The average cell height (basal-to-apical distance in %) (x-axis) and signal intensities (y-axis) were measured and calculated as described in the Materials and Methods section. The shaded areas on each graph represent the areas of the cell (basal 20% and apical 40%) free from influence of the nuclei used for statistical analyses. Number of asterisks refer to the level of statistical significance (n.s. p > 0.05, * p ≤ 0.05, ** p ≤ 0.01, *** p ≤ 0.001, and **** p ≤ 0.0001) between distributions of WT and substituted envelope proteins in the shaded areas. The numbers of midgut enterocytes used to generate the graphs are as follows: VSV G^WT^ n = 69; VSV G^ΔY^ n = 96; GP64^WT^ n = 90; GP64^ΔY^ n = 106. The numbers of salivary gland cells used to generate the graphs are as follows: VSV G^WT^ n = 160; VSV G^ΔY^ n = 211; GP64^WT^ n = 163; GP64^ΔY^ n = 186. Data shown represents combined results from 3 independent experiments. Viral envelope proteins (white) were labelled with mouse anti-VSV G or anti-GP64 primary antibodies and Alexa Fluor 647 donkey anti-mouse secondary antibody. Scale bars represent 5 µm.

### VSV G trafficking in *Drosophila* salivary gland cells differs from that in midgut enterocytes

After exiting infected enterocytes, viruses circulating in the hemolymph infect secondary tissues by interacting with and entering cells via their hemocoel-facing surfaces. The salivary gland is a secondary site of infection that is essential for arbovirus transmission, with an absolute requirement for viral egress into salivary secretions [37]. Thus, the salivary gland epithelium represents a second major tissue barrier to arbovirus transmission. Insect-specific viruses such as baculoviruses may also infect salivary glands, but unlike arboviruses, efficient apical egress of baculovirus progeny virions into saliva is not necessary for their dissemination. Arboviruses, on the other hand, must enter the salivary glands basally but exit apically, an orientation of transit opposite to that in the midgut epithelium (apical entry and basal exit). Viral envelope proteins of arboviruses may therefore traffic differently in the midgut and salivary glands. Because salivary gland epithelial cells have distinctly different cellular architecture and function than midgut enterocytes (Fig. 1A vs. 1C) [10, 17, 38–42], it is unclear whether the same trafficking signals or mechanisms are used in these two barrier tissues. To better understand differences in the cellular architecture of midgut and salivary gland cells, we examined a series of cellular markers using antibodies that identified proteins associated with apical domains (cortical actin), septate junctions (Discs large), and basolateral domains (E-Cadherin) of polarized cells (Fig. 1 and S3). Midgut epithelial cells (enterocytes) are characterized by apical projections that comprise the apical microvilli (Fig. 1A), and a basal region that is a tightly packed labyrinth of small membrane invaginations [43]. In contrast, the polarized salivary gland cells contain large invaginations of the apical surface known as canaliculi that penetrate deep within the cell, and a basal surface that is adjacent to the hemolymph. The apical membrane surfaces of the salivary gland cells line the lumen of the canaliculi, which empty into the salivary duct.

We expressed VSV G in salivary gland cells and analyzed its distribution to determine whether VSV G expressed alone is sufficient for its trafficking to the apical membranes, consistent with the requirements for arbovirus egress. In salivary gland cells, we found that VSV G localized near the apical invaginations in most cells (Fig. 1D, VSV G). The observed patterns of VSV G distribution in salivary gland cells were variable. Patterns ranged from basally enriched (18.8%) to no polarity (52.5%) to apically enriched (28.8%) (Fig. S4, n = 160). Salivary gland cells with distinct apical localization of VSV G are highlighted in Figure 1D, and examples of basal and non-polarized distributions in salivary gland cells are shown in supplemental data (Fig. S4). Thus, VSV G does not have a consistently polarized distribution in salivary glands, which contrasts with the uniformly basal distribution of VSV G in midgut enterocytes. Despite the variability in VSV G distribution, the apical-most 40% region of the salivary gland cells accounted for 49.5 ± 1.14% (mean ± SE, n = 160) of the total VSV G signal (Fig. 2C, VSV G, WT), with the majority (81.3%) of salivary gland cells showing apical presence of VSV G (Fig. S4). The large proportion of cells exhibiting some apical trafficking suggests that this level of apical trafficking could potentially support egress of the virus from salivary gland cells into saliva for transmission.

We similarly examined the distribution of GP64 expressed in adult *Drosophila* salivary gland cells. In contrast to the variable but frequently apical membrane localization of VSV G, GP64 was consistently concentrated in basal regions of salivary gland cells (Fig. 1D, GP64). This basal enrichment of GP64 in the salivary gland cells was similar to that of the midgut enterocytes, with the basal-most 20% region of the salivary gland cells accounting for 56.8 ± 1.53% (mean ± SE, n = 163) of the total GP64 signal (Fig. 2C, GP64, WT). We detected substantially less GP64 in apical membrane invaginations compared with that observed for VSV G (w = 2071, p < 2.2 × 10^-16^), with the apical 40% region of the salivary gland cells accounting for only 22.9 ± 0.95% (n = 163) of the total GP64 signal.

The divergent trafficking of VSV G in midgut enterocytes and salivary gland cells (basal enrichment in midgut enterocytes and more apical localization in salivary gland cells) parallels the anticipated polarized virion budding in these two tissues. The basal localization of GP64 in both tissues is also consistent with the infection cycle of the insect-specific baculovirus, in which viral egress into the hemocoel facilitates dissemination and systemic infection. In both cases, the interaction of each of these viral envelope proteins with the cell trafficking machinery is sufficient for their appropriate basal or apical trafficking, and reflect key differences between insect-specific and insect-vectored virus movement through their insect hosts.

### A YxxØ motif directs polarized trafficking of VSV G and GP64 in *Drosophila* midgut enterocytes

Both VSV G and GP64 are homotrimeric type 1 integral membrane proteins that are functionally and structurally related class III viral fusion proteins [44], despite being from different viral families (ssRNA *Rhabdoviridae* and dsDNA *Baculoviridae*, respectively) with dramatically different infection cycles. Since they are both trafficked to basal membranes when expressed in uninfected midgut enterocytes (Fig. 1B), they may utilize similar strategies or protein motifs to engage insect host cell basal trafficking machinery in the midgut. VSV G and GP64 both contain a canonical tyrosine-based YxxØ motif (Y = tyrosine; x = any amino acid; Ø = a bulky hydrophobic amino acid) in their short cytoplasmic tails (Fig. 2A). The requisite tyrosine (Y) and isoleucine (I) residues of the VSV G YxxØ motif (YCMI) have previously been identified as necessary for its basal trafficking in polarized mammalian cell cultures (MDCK cells) [23, 24]. We therefore hypothesized that the YxxØ motif from VSV G and GP64 might direct basal trafficking of these proteins in *Drosophila* midgut enterocytes *in vivo*. To investigate this, we substituted the Y and I residues in VSV G with alanine (YTDI to aTDa) and incorporated the substituted VSV G construct (VSV G^ΔY^) into a transgenic *Drosophila* line (Fig. 2A, VSV G). We also produced a similar GP64 construct in which the corresponding Y and I residues of the putative YxxØ motif of GP64 were substituted (YCMI to aCMa) and generated a transgenic *Drosophila* line for expressing the modified GP64 (GP64^ΔY^) (Fig. 2A, GP64). The fly lines encoding VSV G or GP64 with alanine substitutions were then used to functionally analyze the role of the YxxØ motif for basal trafficking of both proteins in polarized insect midgut enterocytes.

When VSV G^ΔY^ was expressed in midgut enterocytes, the dramatic basal localization that was previously observed with the VSV G^WT^ protein was lost (Fig. 2B, VSV G, WT vs ΔY). VSV G^WT^ was found concentrated along the basal portion of the cell, suggesting association along the tightly invaginated basal labyrinth, while VSV G^ΔY^ was often adjacent to and above nuclei and in the brush border (apical) (Fig. 2B, Midgut, VSV G, ΔY). Similarly, the dramatic basal enrichment of GP64^WT^ was lost when the modified GP64 (GP64^ΔY^) was expressed (Fig. 2B, Midgut, GP64, WT vs ΔY). To quantify differences in polarized trafficking patterns between WT and modified VSV G and GP64 proteins, we plotted the mean basal-to-apical distributions of the envelope proteins across numerous cells (see Materials and Methods and Fig. S1). Each graph in Figure 2B displays the relative basal-to-apical distribution of the envelope proteins (e.g., VSV G^WT^ vs. VSV G^ΔY^) from 3 separate replicates with approximately 3-5 guts and at least 20 enterocytes per replicate (total n ≥ 69 per genotype). The VSV G^WT^ localization pattern was strongly basal, with approximately 70.4 ± 1.28% of the signal in the basal-most 20% region of the enterocytes (Fig. 2B, VSV G, WT, open circles). In contrast, VSV G ^ΔY^ was significantly less basally enriched with only 32.5 ± 1.20% of the signal in the basal-most 20% of the enterocytes (t = 21.6, df = 161, p < 2.2 × 10^-16^) (Fig. 2B, VSV G, ΔY, filled squares). When the same comparison was performed with GP64^WT^ and GP64^ΔY^, a similar loss of basal enrichment of GP64^ΔY^ (37.5 ± 1.13% basal signal) was detected when compared to GP64^WT^ (70.3 ± 1.25% basal signal) (t = 19.4, df = 188, p < 2.2 × 10^-16^) (Fig. 2B, GP64, WT vs ΔY). GP64^ΔY^ signals were more diffuse within cells compared to GP64^WT^, and GP64^ΔY^ was sometimes detected in the apical brush border. These data indicate that the YxxØ motif in the cytoplasmic tails of both VSV G and GP64 are functional motifs required for the basal trafficking of each viral envelope protein in polarized insect midgut enterocytes.

### The YxxØ motif is not a prime determinant of polarized trafficking of VSV G or GP64 in *Drosophila* salivary gland cells

Our prior analysis of protein markers of cell polarity showed that the apical-basal orientation of midgut enterocytes and salivary gland cells were similar, with basal membranes adjacent to the hemocoel and the apical membranes bordering the lumen of the organ. To determine whether alteration of the YxxØ motif would impact VSV G trafficking in salivary gland cells, we compared the distribution patterns of VSV G^WT^ and VSV G^ΔY^. Tyrosine motif alteration resulted in slightly reduced basal VSV G localization, with the basal-most 20% region of the salivary gland cells accounting for 23.4 ± 0.97% of the total VSV G^WT^ signal and 13.8 ± 0.56% of the total VSV G^ΔY^ signal (w = 25056, p = 1.33 × 10^-15^), but did not change the overall distribution pattern and only slightly increased VSV G localization in the apical-most 40% region of the salivary gland cells (49.5 ± 1.13% to 53.2 ± 0.79%, t = −2.62, df = 298, p = 9.25 × 10^-3^) (Fig. 2C, VSV G). In salivary glands expressing VSV G^ΔY^, we found that the proportion of cells with apically localized VSV G^ΔY^ (95.7%, n = 211) was slightly increased compared to cells expressing VSV G^WT^ (81.3%, n = 160) (χ^2^ = 18.8, df = 1, p = 1.46 × 10^-5^) (Fig. S4). Also, the proportion of cells with basally localized VSV G^ΔY^ was slightly decreased in comparison to VSV G^WT^ (13.8 ± 0.56% for VSV G^ΔY^ vs. 23.4 ± 0.97% for VSV G^WT^, w = 25056, p = 1.33 × 10^-15^) (Fig. S4). Thus, ablation of the YxxØ motif did not result in a substantial change in the overall distribution pattern (Fig. 2C, VSV G, graph; and Fig. S4), suggesting limited involvement of the YxxØ motif in the apical trafficking of VSV G in salivary gland cells. It is important to note that since apical membranes of salivary gland cells extend deep into the cytoplasm (Fig. 1C), our method for quantifying basal-to-apical distribution in salivary gland cells may underestimate apical enrichment of VSV G (i.e., VSV G decorating the apical membrane reaching the level of the nucleus would not appear apical in the graphs). Nevertheless, our results indicate that VSV G is apically localized in most salivary gland cells, and the VSV G YxxØ motif appears to play only a minimal role in its apical trafficking in *Drosophila* salivary gland cells.

In contrast to the apical distribution of VSV G^WT^ in the salivary gland cells, GP64^WT^ was concentrated at basal membranes in salivary gland cells (Fig. 2C, GP64). This localization of GP64 was similar to that observed in midgut enterocytes (Fig. 1D vs. 2C, GP64). The basal enrichment of GP64^ΔY^ (Fig. 2C, GP64, ΔY) was only marginally reduced in the basal-most 20% region of the salivary gland cells when compared to GP64^WT^ (52.3 ± 1.55% for GP64^ΔY^ vs 56.8 ± 1.53% for GP64^WT^, w = 17017, p = 0.048) with no influence on the overall distribution pattern. This suggests minimal involvement of the YxxØ motif in the basal trafficking of GP64 in the salivary gland cells.

Altogether, our data suggest that while the YxxØ motifs are required for basal trafficking in midgut enterocytes, they are mostly dispensable for directing trafficking of either GP64 or VSV G in salivary gland cells. Furthermore, the observation that VSV G is found localized to apical membranes in salivary gland cells while strictly basal in enterocytes, whereas GP64 is strictly basal in both tissues, indicates that there are additional, yet unidentified trafficking/sorting motifs in VSV G and GP64, as well as differences in the mechanisms of polarized trafficking between midgut enterocytes and salivary gland epithelial cells. Both aspects are topics of interest for future studies.

### Clathrin AP complexes 1 and 3 direct basal trafficking of VSV G in *Drosophila* midgut enterocytes

We next aimed to identify key host factors involved in the basal trafficking of viral envelope proteins in midgut enterocytes. Clathrin adaptor protein (AP) complexes direct a large variety of membrane trafficking events in the cell. This may include trafficking among compartments (ER, Golgi, endosomes, lysosomes, etc.) as well as polarized and non-polarized trafficking to subdomains of the plasma membrane. In models of polarized trafficking of membrane proteins in mammalian cells, interactions between membrane protein YxxØ motifs and the μ subunits of clathrin AP complexes are early steps in protein trafficking to basal membranes. Also, in a prior RNAi screen of cultured *Drosophila* cells [25], we found that RNAi of clathrin (*chc*) and AP genes (*AP-2u* and *AP-1-2β*) resulted in decreased GP64 transport to the cell surface. Although AP-1-2β is not expected to directly interact with the YxxØ motif, depletion of any one of the four canonical subunits of AP complexes is expected to render its corresponding complex nonfunctional. In Drosophila and other insects, AP-1-2β is shared between the heterotetrameric AP-1 and AP-2 complexes, both of which would therefore be inactivated upon AP-1,2β knockdown. Since we identified the importance of the YxxØ motif in the basal trafficking of VSV G in midgut enterocytes, we next examined the roles of the three AP complexes in the polarized transport of VSV G in that tissue.

To probe the role of each AP complex, a fly line (*Myo-Gal4, UAS-nlsGFP, tub-Gal80^ts^; UAS-VSV G^WT^*) with temperature-regulated midgut expression of WT VSV G, was crossed with RNAi lines to obtain F1 progeny co-expressing VSV G and an RNAi construct targeting (separately) each one of three clathrin adaptor complexes. We then compared VSV G distribution patterns in each of the AP complex knockdowns, to negative controls (midguts from driver/*AttP2* background control, lacking any RNAi). VSV G showed strong basal enrichment in enterocytes of control midguts (Fig. 3, Control, top panels). However, when VSV G was co-expressed with RNAi constructs that targeted AP-1µ or AP-3µ, the strong basal localization of VSV G was lost, and VSV G was distributed less basally (32.3 ± 0.771% and 45.2 ± 1.39%, respectively) compared to the control (56.9 ± 1.06%) in the basal 20% region of the enterocytes (Pairwise Wilcoxon rank sum test: p < 2.00 × 10^-16^ and p = 6.00 × 10^-9^, respectively) (Fig. 3, Control vs. AP-1µ and AP-3µ). This contrasts with the result from the knockdown of AP-2µ (53.2 ± 1.38%), in which VSV G distribution appeared similar to that of control enterocytes (56.9 ± 1.06% basal) (Pairwise Wilcoxon rank sum test: p = 0.126) (Fig. 3, Control vs. AP-2µ). The patterns observed in the presence of AP RNAi targeting AP-1µ or AP-3µ were similar to that observed when the YxxØ motif was ablated in construct VSV G^ΔY^ (Fig. 2B, VSV G, ΔY). These patterns were consistent across independent guts (≥3) and individual cells (N ≥ 119) after 3 days of RNAi. We also observed a similar disruption of basal trafficking from an RNAi line targeting AP-1,2β (40.4 ± 0.958%), which served as a positive control (Pairwise Wilcoxon rank sum test: p < 2.00 × 10^-16^) (Fig. 3, Control vs. AP-1,2β). In addition to the above results, pharmacological disruption of the YxxØ-AP interactions using anthranilic acid (ACA) [45] reduced VSV G^WT^ localization in the basal-most 20% region of midgut enterocytes (65.4 ± 1.97% in control vs. 50.4 ± 1.53% in ACA treated; w = 4154, p = 3.48 × 10^-8^) (Fig. S5). Thus, our results from RNAi knockdowns of AP complexes combined with results from YxxØ ablation in VSV G, suggest that AP-1 and AP-3 but not AP-2 complexes are necessary for YxxØ-mediated VSV G basal trafficking in midgut enterocytes.

**Figure 3.**
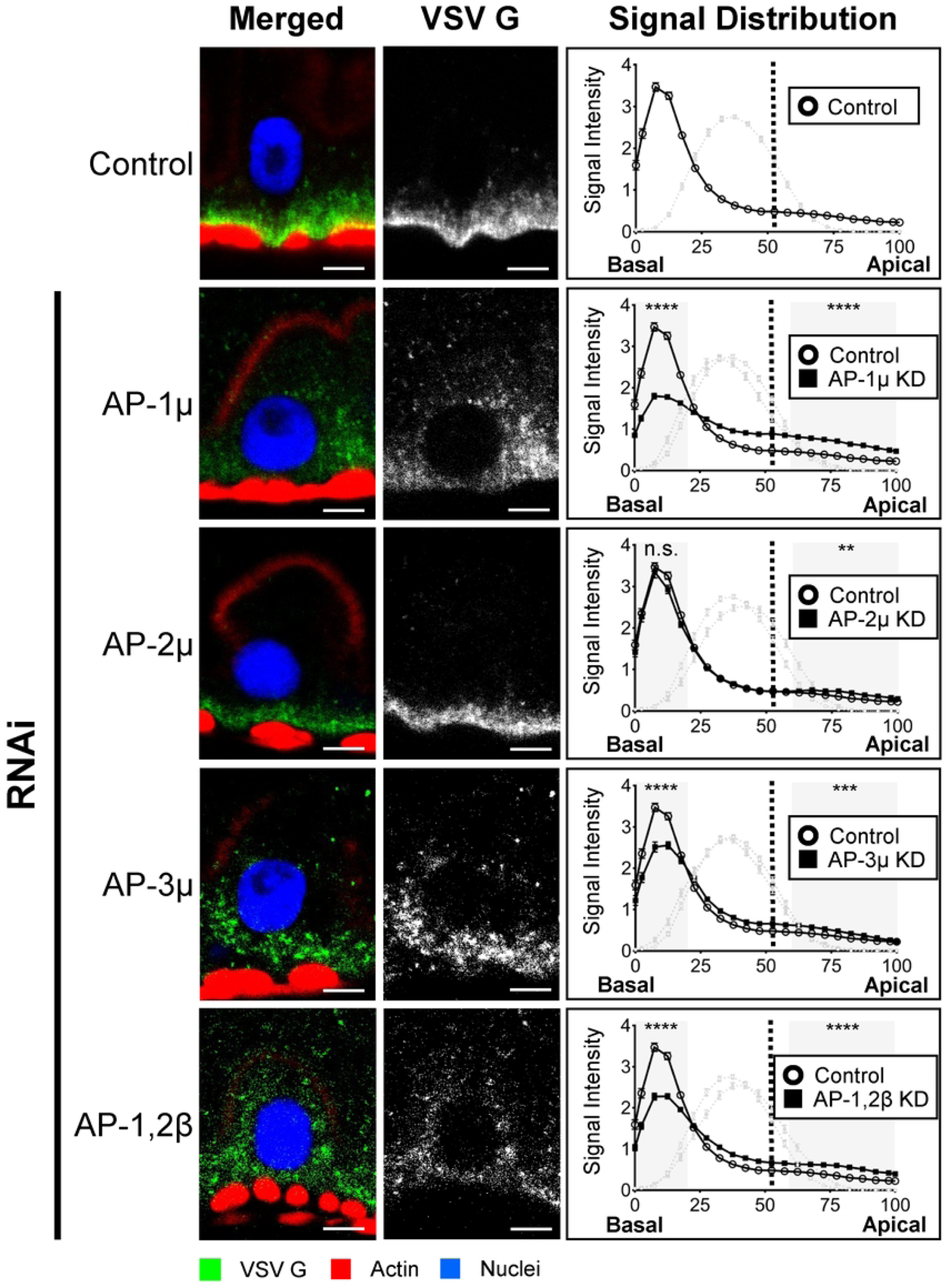
Effects of RNAi knockdown of clathrin adaptor protein subunits on basal trafficking of VSV G in *Drosophila* midgut enterocytes. The image panels (left) show the localization of VSV G (green in images on the left and white in images on the right) in enterocytes from either control flies (top row, no knockdown) or from flies co-expressing VSV G and RNAi constructs to deplete AP-1µ, AP-2µ, AP-3µ, or AP-1,2β (indicated on the left of each row). The graph in each row compares the basal-to-apical distribution of VSV G in control enterocytes (top panels) against enterocytes from flies co-expressing VSV G and an RNAi construct (indicated on the left of each row of image panels). RNAi of AP-1µ, AP-3µ, and AP-1,2β resulted in reduced basal enrichment of VSV-G in enterocytes. For all knockdowns, distribution of VSV G was analyzed after 3 days of RNAi induction under the midgut *Myo-Gal4 tub-Gal80^ts^* driver. The average relative position of the nucleus was also plotted as a gray line. The average cell height (basal-to-apical distance in %) (x-axis) and VSV G signal intensities (y-axis) were measured and calculated as described in the Materials and Methods section. The basal and apical regions on the graphs were established based on the location of the septate junction (discs-large polarity marker) (Fig. S1), and the division is indicated at 52.5% on the x-axis by a vertical dashed line. The shaded areas on each graph represent the areas of the cell (basal 20% and apical 40%) relatively free from influence of the nuclei and used for statistical analyses. The number of asterisks in the shaded areas refer to the level of statistical significance (n.s. p > 0.05, * p ≤ 0.05, ** p ≤ 0.01, *** p ≤ 0.001, and **** p ≤ 0.0001) in VSV G distribution differences between RNAi and control flies, in those areas. The numbers of midgut enterocytes analyzed were as follows: Control (n = 165), AP-1µ RNAi (n = 141), AP-2µ RNAi (n = 119), AP-3µ RNAi (n = 122), and AP-1,2β RNAi (n = 146). Data shown represents combined results from 3 independent experiments. Viral envelope protein (green) was labelled with mouse anti-VSV G primary antibody and Alexa Fluor 647 donkey anti-mouse secondary antibody, actin (red) was labelled with Alexa Fluor 555 phalloidin, and nuclei (blue) were stained with DAPI. Scale bars represent 5 µm.

### Specific Rab GTPases are important for basal trafficking of VSV G in *Drosophila* midgut enterocytes

Rab GTPases are known to be master regulators that direct vesicular protein sorting and trafficking. Their numerous roles include the regulation of vesicle budding, transport, tethering, and fusion with target membranes [46]. Specific Rab GTPases may also serve different roles in different cell types [47]. Importantly, they have also been shown to be required for basal trafficking of host and viral proteins [48–54]. Rab GTPases are engaged in the establishment and maintenance of cell polarization [55, 56] and the specific subcellular localization of some Rab GTPases may differ in different cell types or tissues. We aimed to identify Rab GTPases that could be involved in the basal trafficking of VSV G in enterocytes. Our approach was to first select two groups of candidate Rab genes: (1) Rab genes that have previously been shown to be important for viral protein trafficking in other cellular systems and (2) Rab genes that showed interesting apical-basal localization in enterocytes, and that have different localization patterns in enterocytes compared to salivary gland cells. For the first group, we had previously identified Rab1 and 4 as essential for the transport of GP64 to the surface of non-polarized cultured *Drosophila* cells [25]. Although we did not identify Rab11 as important in that study, we included Rab11 in the current study due to its involvement in sorting/recycling endosome function, similar to Rab4, which was important for GP64 surface trafficking [25, 57]. It was previously shown that basal trafficking of VSV G in mammalian MDCK cells requires coordination of the AP-1 complex with Rab8 [49] and that Rab10 is required for basal secretion of *Drosophila* gut basal lamina components [53]. Therefore, we investigated whether Rab1, 4, 8, 10, and 11 play any roles in basal VSV G trafficking in insect midgut enterocytes. To select a second group of candidate Rab GTPases, we performed a comprehensive comparative analysis of the localization patterns of 30 YFP-tagged Rab GTPases [58] in *Drosophila* midgut enterocytes and salivary gland cells (Fig. S6). For this analysis, UAS-driven YFP-tagged Rab GTPases were ubiquitously expressed under the *da-Gal4 tub-Gal80^TS^* driver. and we compared the distribution of each YFP-tagged Rab GTPase in midgut vs salivary gland cells by confocal microscopy. We identified several YFP-Rab GTPases with basal enrichment (Rab8, 10, and 30) or both basal and apical enrichment (Rab23 and 35) in enterocytes, and with salivary gland cell localization patterns that differed from that of midgut enterocytes, displaying either apical enrichment in the canaliculi (Rab23 and 35) or a non-polarized localization pattern (Rab8, 10, and 30), similar to the localization patterns observed for VSV G in each cell type (Fig. 4).

**Figure 4.**
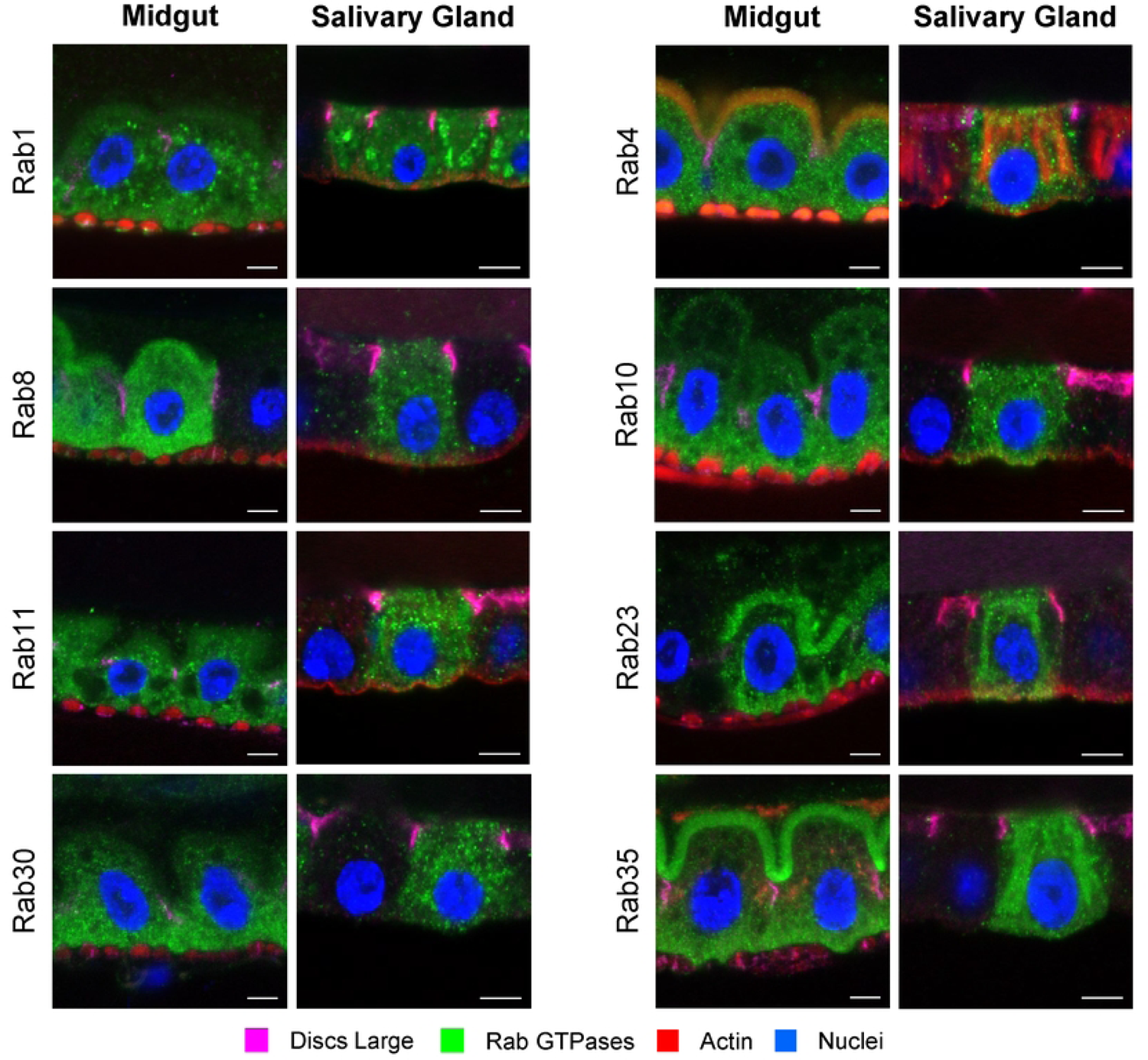
Rab GTPase localization patterns in midgut enterocytes and salivary gland cells. Representative images of immunostained YFP-Rab GTPases (green) illustrate differences in localization patterns in *Drosophila* midgut enterocytes and salivary gland cells. Rab8, 10, 23 and 30 showed basally enriched localization patterns in midgut enterocytes, but showed non-polarized or more apically-enriched localization patterns in salivary gland cells, suggesting that they may be involved in the differential trafficking of VSV G in the two tissues. The YFP-tagged Rab GTPases were expressed ubiquitously in adult *Drosophila* under the *da-Gal4 tub-Gal80^ts^* driver for 3 days before dissection and immunostaining with chicken anti-GFP primary antibody and Alexa Fluor 488 goat anti-chicken secondary antibody (green). Actin (red) was labelled with Alexa Fluor 555 phalloidin, nuclei (blue) were stained with DAPI, and Discs large (magenta) was stained with mouse anti-Discs large primary antibody and Alexa Fluor 647 donkey anti-mouse secondary antibody. Scale bars represent 5 µm.

Next, we examined these selected Rab GTPases to assess their potential roles in basal trafficking of VSV G in *Drosophila* midgut enterocytes. We co-expressed VSV G with RNAi constructs that separately targeted each of the following Rab GTPases: Rab1, 4, 8, 10, 11, 23, 30, or 35 (Fig. 5) by crossing with a fly line with temperature-regulated VSV G midgut expression (*Myo-Gal4, UAS-nlsGFP, tub-Gal80^ts^; UAS-VSV G^WT^*). F1 progeny flies were shifted to the permissive temperature to allow VSV G/Rab RNAi co-expression for 5 days (with the exception of Rab11 RNAi, which was co-expressed with VSV G for 3 days) before dissecting and immunostaining midgut enterocytes to assess VSV G localization. Basal enrichment of VSV G was significantly reduced in midgut enterocytes of flies co-expressing VSV G and each of the RNAi constructs targeting: Rab1 (42.2 ± 0.889%), Rab4 (42.8 ± 0.705%), Rab8 (36.6 ± 0.884%), Rab10 (39.1 ± 1.16%), Rab23 (41.8 ± 0.884%), Rab30 (41.6 ± 1.07%), and Rab35 (44.7 ± 1.08%) compared to their respective controls (Attp2 for Rab1, 4, 8, and 10: 53.7 ± 0.980%; Attp40 for Rab23 and 35: 56.3 ± 0.839%; and Attp2 for Rab30: 56.9 ± 1.06%) (Pairwise Wilcoxon rank sum test: p = 3.9 × 10^-14^, p = 3.4 × 10^-14^, p < 2.0 × 10^-16^, p = 5.2 × 10^-16^, p < 2.0 × 10^-16^, and p = 1.3 × 10^-14^, respectively for each Rab RNAi compared to its control), (Fig. 5, Control vs. RNAi Rab#). Surprisingly, we found no substantial effect on VSV G basal localization from the Rab11 knockdown (79.1 ± 1.12%) compared to the control (78.0 ± 1.32%) (w = 7547, p = 0.665) (Fig. 5, Control vs. RNAi Rab11). Thus, our analysis revealed that all but one of the Rab GTPases selected for analysis (Rab1, 4, 8, 10, 23, 30 and 35) were involved in basal trafficking of VSV G in midgut enterocytes.

**Figure 5.**
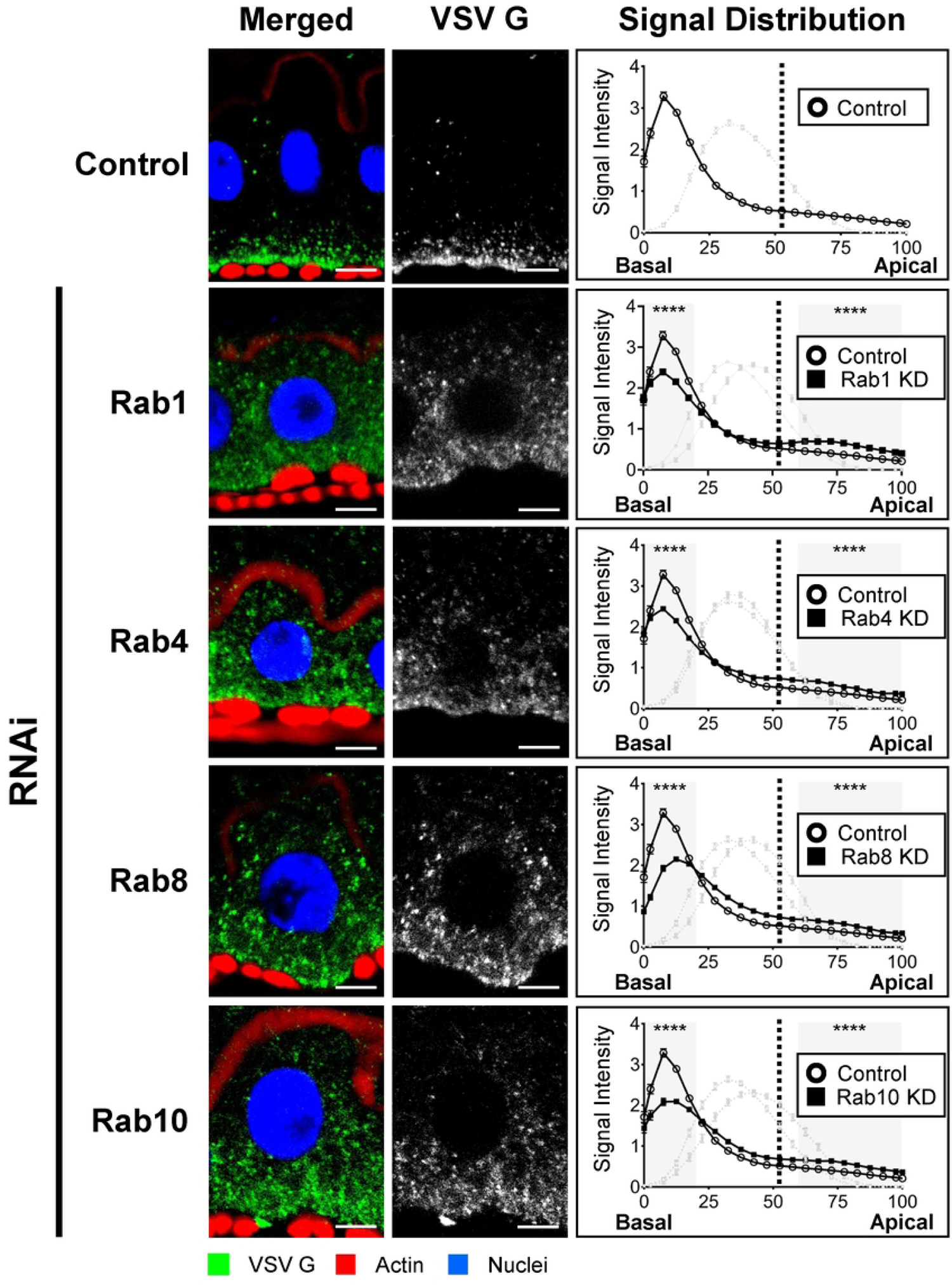
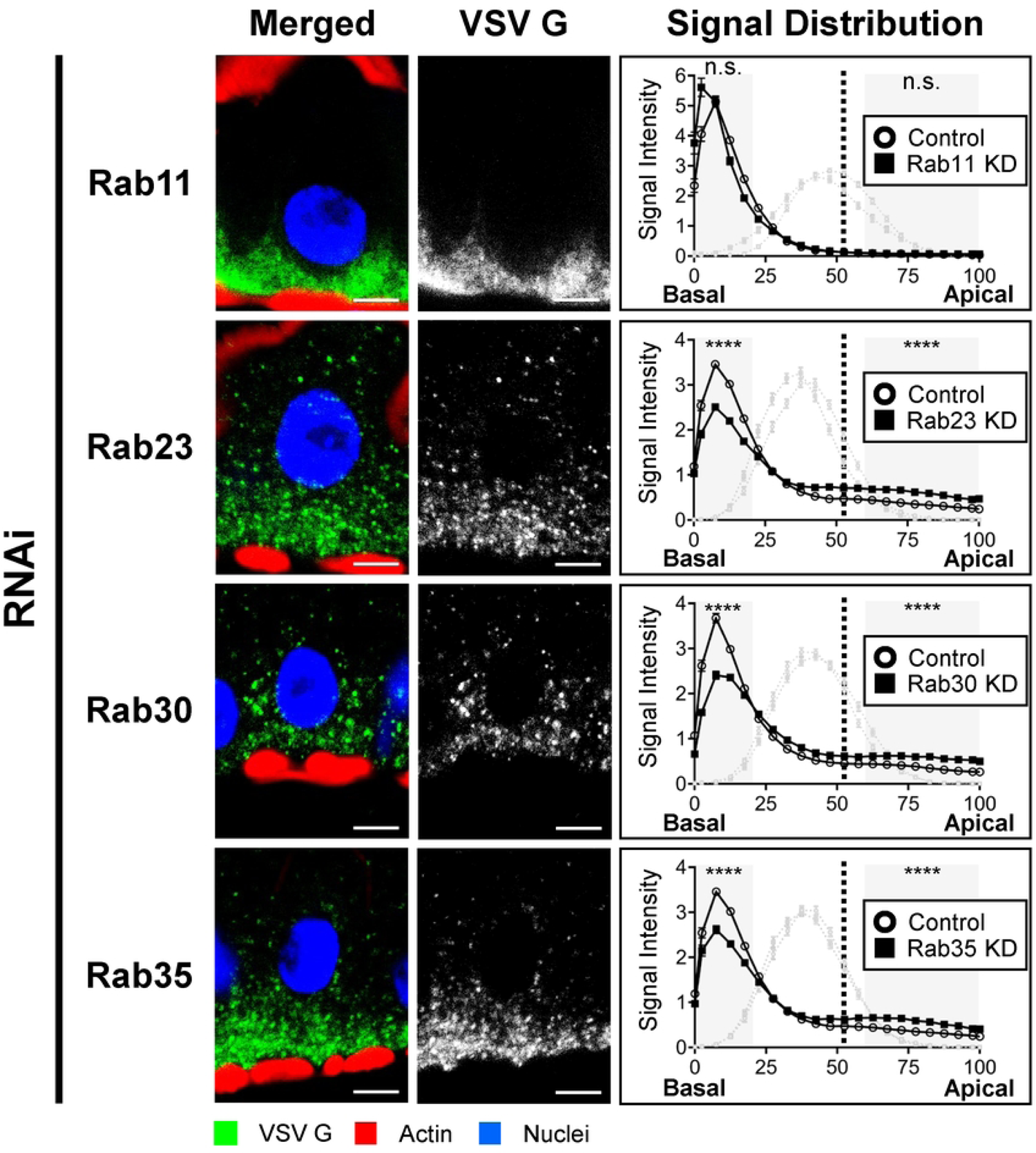
Effects of RNAi knockdowns of selected Rab GTPases on basal trafficking of VSV G in *Drosophila* midgut enterocytes. The image panels (left) show the localization of VSV G (green in images on the left and white in images on the right) in enterocytes from either control flies (top row, no knockdown) or from flies co-expressing VSV G and RNAi constructs to deplete Rab1, Rab4, Rab8, Rab10, Rab11, Rab23, Rab30, or Rab35 (indicated on the left of each row of image panels). RNAi of all selected Rab GTPases (except Rab11) resulted in reduced basal enrichment of VSV-G in enterocytes. For all knockdowns (except Rab11), distribution of VSV G was analyzed after 5 days of RNAi induction (3 days for Rab11) under the midgut *Myo-Gal4 tub-Gal80^ts^* driver. The average relative position of the nucleus was also plotted as a gray line. The average cell height (basal-to-apical distance in %) (x-axis) and VSV G signal intensities (y-axis) were measured and calculated as described in the Materials and Methods section. The basal and apical regions on the graphs were established based on the location of the septate junction (discs-large polarity marker) (Fig. S1), and the division is indicated at 52.5% on the x-axis by a vertical dashed line. The shaded areas on each graph represent the areas of the cell (basal 20% and apical 40%) relatively free from influence of the nuclei and used for statistical analyses. The number of asterisks in the shaded areas of the graphs refer to the level of statistical significance (n.s. p > 0.05, * p ≤ 0.05, ** p ≤ 0.01, *** p ≤ 0.001, and **** p ≤ 0.0001) in VSV G distribution between RNAi and control flies. The numbers of midgut enterocytes analyzed were as follows: Control (n = 572), Rab1 RNAi (n = 140), Rab4 RNAi (n = 150), Rab8 RNAi (n = 138), Rab10 RNAi (n = 124), Rab11 (n = 131), Rab23 RNAi (n = 210), Rab30 RNAi (n = 207), Rab35 RNAi (n = 235). Data shown represents combined results from 3 independent experiments with each knockdown compared to its appropriate control. Viral envelope protein (green) was labelled with mouse anti-VSV G primary antibody and Alexa Fluor 647 donkey anti-mouse secondary antibody, actin (red) was labelled with Alexa Fluor 555 phalloidin, and nuclei (blue) were stained with DAPI. Scale bars represent 5 µm.

## Discussion

Viral trafficking within insect hosts is critical for the success of the viral life cycle. Because little is known about how insect viruses navigate across important tissue barriers in their insect hosts, we examined the subcellular trafficking of two model viral envelope proteins in two critical barrier tissues: midgut and salivary glands. In addition to their roles as barriers and bottlenecks to virus infection and transmission, midgut and salivary glands serve distinct physiological roles in insects. The midgut is the primary tissue for absorption of nutrients and their transfer to the circulatory system. As such, the protein trafficking pathways in the midgut are specialized for apical secretion of digestive enzymes and components of the peritrophic matrix, as well as for absorption of nutrients from the gut lumen. Absorbed nutrients are transported to the basal surfaces of midgut enterocytes, where they are delivered to the hemocoel. In addition, the basal lamina, a thick collagen-containing matrix, is secreted from the basal surfaces of midgut enterocytes. Thus, protein transport and secretory pathways within midgut enterocytes perform many central functions that require precise recognition of proteins and targeted polarized transport. Insect salivary glands are thought to primarily produce secretions that play roles in feeding, particularly in food processing (pre-digestion, lubrication) and sometimes in manipulation of the host during blood feeding (anticoagulation, anesthesia, vasodilation, etc.). Because food may be acquired in large quantities during a short period, salivary glands employ mechanisms for producing then transporting and secreting salivary proteins (and other molecules) from apical membranes, and delivering them on demand. In some insects, such as mosquitoes, salivary gland cells appear to secrete large quantities of saliva and store them in large apical cavities (acinar cavities) [17, 38, 40, 59]. In other insects, such as blow flies, the apical membrane of the salivary gland cells forms deep invaginations (canaliculi) that greatly increase membrane surface area for secretion [60]. As such, the salivary glands represent a robust secretory and release apparatus, but the cellular architecture (invaginated apical membranes) and cell types (the lack of visceral muscles and stem cells) differ fundamentally from that of the midgut. Many viruses that infect insects have adapted to utilize the complex transport systems in various cell types to move infection through the animal successfully. Such navigation of viral infection through the animal is particularly interesting in the case of arboviruses, but it is also crucial for the success of insect-pathogenic viruses.

To examine viral envelope protein trafficking through these polarized tissues of insects, we used transgenic *Drosophila* to express genes encoding either baculovirus (AcMNPV) GP64 or an arbovirus (VSV) G in these tissues. Upon expression, each viral envelope protein was transported to and concentrated at the basal membranes of midgut enterocytes (Fig. 1) in the absence of infection or aid of any other viral proteins. This result mirrors the basal envelope protein trafficking reported during infections (VSV-infected MDCK cells and AcMNPV-infected *T. ni* midgut) [23, 24, 27], and demonstrates that the information required for polarized envelope protein trafficking in insect midgut enterocytes is encoded within the GP64 and VSV G proteins.

We also examined polarized trafficking of these proteins in salivary gland cells. Salivary gland cells are specialized for secreting proteins into the salivary ducts. Arboviruses utilize salivary gland trafficking pathways to exit the insect vector host and facilitate infection of the vertebrate host during blood feeding. Because this viral transit appears to be in an orientation that is opposite to that observed in midgut enterocytes, we first used a series of polarity markers to determine the apical-basal orientation of cells within the salivary gland epithelium (Fig. S3). Indeed, we confirmed that apical membrane domains face the lumen of the salivary glands, and basal membranes are adjacent to the hemocoel. We show that GP64 was consistently localized to basal membranes of salivary gland cells, similar to the results from midgut enterocytes. In contrast, yet consistent with the arbovirus life cycle, we found that VSV G was trafficked to apical membranes that line the apical invaginations of salivary gland cells. While VSV G trafficking was not strictly apical in all cells, we found that 28.8% of SG cells had distinctly apical trafficking, while another 52.5% of cells had non-polarized trafficking, and thus some apical presence of VSV G. This suggests apical budding was possible from approximately 81% of SG cells. The observed partial or inconsistent polarized trafficking may indicate that viral infection could provide additional factors necessary for robust apical trafficking of G in salivary gland cells. It is also possible that partial apical trafficking is sufficient for VSV transmission, and basal trafficking in salivary gland cells may also play a role in the infection cycle. It is of note that some alphaviruses have been observed to bud into both acinar cavities and from lateral and basal membranes of infected mosquito salivary gland cells [38, 59]. While we have focused on a powerful model insect for the current studies, it will be of great interest to examine viral envelope protein trafficking in mosquitoes and other natural hosts since they may have different trafficking mechanisms for arboviruses than *Drosophila*. Most importantly, our results show that when GP64 or VSV G was expressed alone, the polarized trafficking of each protein was sufficient to reach membrane locations appropriate to the infection cycles of their parent viruses. These results from *Drosophila* tissues also suggest that the polarized trafficking pathways in midgut and salivary gland cells are widely conserved in insects and that the signals for appropriate polarized trafficking are largely if not entirely encoded in the protein sequences or structures of these viral envelope proteins.

In prior studies of VSV G trafficking in mammalian MDCK cells, a YxxØ (YTDI) amino acid sequence motif in the cytoplasmic tail was identified as necessary for its basolateral targeting [23, 24]. When examined in insect midgut enterocytes, we found that alanine substitutions of the key residues in the YxxØ motif resulted in disruption of polarized trafficking, and we concluded that the YxxØ motif is necessary for basal trafficking of VSV G in *Drosophila* midgut enterocytes (Fig. 2). In parallel experiments, we identified a similar YxxØ (YCMI) motif in the GP64 cytoplasmic tail and found that it was also necessary for basal trafficking. Thus, for both VSV G and baculovirus GP64, the YxxØ motif appears to play a critical role in directing polarized trafficking in *Drosophila* midgut enterocytes (Fig. 2B). Surprisingly, when the same modified VSV G and GP64 constructs (containing an ablated YxxØ motif) were examined in salivary gland cells, we observed no substantial effect on polarized trafficking there (Fig. 2C). While the YxxØ motif is necessary in the midgut epithelium, other signals or motifs appear to be required for polarized trafficking in the salivary gland cells. Thus, the pathways or mechanisms involved in VSV G trafficking appear to differ in the midgut enterocytes and salivary gland cells.

In prior studies of cultured mammalian cells, the YxxØ motif was shown to interact with clathrin adapter protein (AP) complexes to direct basal membrane targeting of proteins. [61–65]. In mammals and plants, there are five different AP complexes, whereas in *C. elegans,* yeast, and *Drosophila,* there are three [66]. We therefore examined the role of each of the 3 AP complexes from *Drosophila* in basal trafficking of VSV G in insect enterocytes. Using RNAi knockdowns of the μ subunit of each complex (the subunit that binds the YxxØ motif), we found that disruption of either the AP-1 or AP-3 complex resulted in disruption of basal trafficking of VSV G in midgut enterocytes (Fig. 3). Furthermore, knockdown of AP-1,2β (which is a beta adaptin shared by the AP-1 and AP-2 heterotetrameric complexes in *Drosophila* and possibly many other lower eukaryotes [66]), resulted in disrupted VSV G basal trafficking. Importantly, the effects of these specific AP complex knockdowns were similar to the effects observed from the ablation of the YxxØ motif in the VSV G protein (Fig. 2B vs. 3).

The AP-1 complex localizes to the trans-Golgi Network (TGN) and recycling/sorting endosomes and is known to direct clathrin-coated vesicular traffic bidirectionally between these compartments and to basolateral plasma membranes [67]. Similar to AP-1, the AP-3 complex is also found on endosomal membranes and the TGN, but in discrete sites that are not overlapping with AP-1. AP-3 has been reported to be involved in cargo transport from early to late endosomes, in the production of lysosome-related organelles in epithelia [61], and in the release of exosomes from neurons [68]. The basal trafficking of VSV G in midgut epithelium was not substantially affected when the AP-2 complex was disrupted (Fig. 3), consistent with the typical role of the clathrin AP-2 complex in directing endocytosed cargo inward (retrograde) from the plasma membrane. Although VSV G, like other YxxØ-motif-containing proteins [69–71], has the potential to be recycled from the plasma membrane to internal organelle membranes, we did not observe an impact on VSV G basal trafficking in enterocytes when the AP-2 complex was disrupted.

Rab GTPases are key regulators of vesicular trafficking, and we therefore examined selected Rab GTPases for their roles or requirements in basal trafficking of VSV G in insect midgut enterocytes. To better understand how the trafficking pathways might differ in midgut enterocytes and salivary gland cells, and to select Rab GTPases for genetic analysis, we used a bank of fly lines expressing Rab-YFP constructs to map and compare Rab localization in midgut enterocytes and salivary gland cells. We identified eight Rab GTPases (Rab1, 4, 8, 10, 11, 23, 30, and 35) that were either differentially localized in cells of the two tissues (Fig. 4 and S6) and/or predicted to be involved in basal trafficking from studies in other systems. We selected these eight Rab genes for analysis by RNAi knockdowns. We found that knockdowns of Rab1, 4, 8, 10, 23, 30, and 35 resulted in substantial reductions in basal localization of VSV G compared to the control (Fig. 5). In the case of the RNAi knockdown of Rab11, VSV G localization was similar to that of the control (Fig. 5; Control vs Rab 11-KD).

Certain Rab GTPases (Rabs4, 8, 10) identified here have been previously associated with basolateral trafficking and/or coordination with clathrin AP complexes [72]. Rab4 is a component of the sorting/recycling endosome, and VSV G has been reported in some mammalian cells to require recycling endosomes for its delivery to the plasma membrane [73]. In midgut enterocytes, it is unclear whether Rab4 is required for the direct basolateral trafficking of G to the plasma membrane, recycling of VSV G, or both. Rab4 and 11 have been identified as regulating distinct “fast” and “slow” recycling pathways, respectively [74]. Because the Rab11 knockdown did not appear to affect VSV G basal trafficking, we conclude that if recycling is involved in basal trafficking, it is likely due to a Rab4-dependent slow pathway, and the Rab11-based pathway may not play a substantial role. Rab8 depletion has been shown to affect basolateral trafficking of VSV G in mammalian MDCK cells [49]. When Rab8 was depleted, it was reported that only basolateral trafficking of VSV G through the secretory pathway (and not through a recycling pathway) was disrupted [48]. When we depleted Rab8 in insect midgut enterocytes, VSV G basolateral trafficking was disrupted, producing a reduced basal VSV G distribution like that observed from Rab1 and Rab4 depletion. This is consistent with results from another study that found Rab8 to be critical for basolateral trafficking of basal lamina (BL) constituents (collagen IV) in polarized *Drosophila* follicular epithelium. In addition, RNAi targeting Rab10 disrupted basal enrichment of VSV G. Rab10 was also previously shown to aid in directing basolateral trafficking of *Drosophila* follicular basal lamina components [53]. Thus, several Rab GTPases identified in our localization mapping and RNAi knockdowns have been identified as important for basal trafficking of host cell proteins in other epithelial cell types.

In the case of most insect pathogenic viruses, such as baculoviruses, the secondary infection of many tissues leads to the death of the insect, and progeny virions are liberated from the insect carcass. For arthropod-vectored viruses, progeny virions produced during secondary infection of salivary glands must be released apically into the salivary duct to facilitate transmission to vertebrate hosts. This divergent scenario of the apical release of virions from salivary glands, relative to the basal virion release from enterocytes, was reflected by apical (or non-polarized) trafficking of VSV G in salivary glands and basal trafficking of VSV G in insect midgut enterocytes. Although VSV G was enriched at the basal-most compartment of some salivary gland cells, a substantial percentage of cells exhibited strongly apical VSV G enrichment in the canaliculi of salivary gland cells. This indicates that apical VSV G trafficking occurs in these cells in the absence of viral infection. This divergent midgut vs. salivary gland VSV G trafficking pattern, combined with differences in Rab GTPase localization patterns in the two tissues, provides substantial evidence that the composition or function of trafficking machinery differs in midgut enterocytes and salivary gland epithelial cells. Interestingly, GP64 expressed in salivary gland cells was not trafficked to apical membrane surfaces (as observed for VSV G) but localized basally as observed in midgut enterocytes (Fig. 1B and D; VSV G vs. GP64). Polarized budding of baculovirus virions has not been studied in insect salivary glands, but our observation of GP64 trafficking in *Drosophila* salivary glands suggest that baculovirus virions may be released from basal membranes of the salivary glands to supplement the systemic infection in these insect-specific pathogens. It is also of note that mutation of the YxxØ motif in both VSV G (G^ΔY^) and GP64 (GP64^ΔY^) did not substantially affect the polarized trafficking pattern of either protein in salivary gland epithelium, in contrast to the effects observed in midgut enterocytes (Fig. 2B, C). Combined with observed differences in Rab localization patterns in midgut vs. salivary gland cells, this indicates that insect salivary gland cells may have a functionally different membrane protein trafficking network compared to insect midgut enterocytes. The observation that WT VSV G and GP64 traffic in opposite directions in salivary gland cells, while both were minimally affected by mutation of the YxxØ motif in salivary glands, also indicates that there are likely divergent but as yet unidentified trafficking signals embedded within the VSV G and GP64 sequences or structures.

In the current studies, we examined two critical issues in virus interactions with their insect hosts: 1) how viruses (insect-specific and insect-vectored) traffic through the polarized midgut epithelium, the first cellular barrier, to establish systemic infection in the insect host; and 2) how arboviruses move across a second polarized epithelial cell barrier (salivary gland cells) in an opposite direction. To address these issues, we developed and used a powerful genetic system to examine: a) viral envelope protein trafficking in each of these tissues, b) viral protein encoded signals for directing polarized trafficking in both tissues, and c) host cell factors necessary for mediating the polarized trafficking of these viral envelope proteins in midgut cells. Using the power of *Drosophila* genetics combined with two well-studied viral envelope proteins, we found that VSV G and GP64 each contain the encoded signals required for basal trafficking in the midgut epithelium, a necessity for virus budding into the hemocoel and establishing systemic infection, and in the case of VSV G, apical trafficking in the salivary gland cells, a route for virus egress into the saliva to be transmitted, consistent with the life cycle of both viruses. We found that the YxxØ motif was necessary for basal trafficking of both proteins in midgut enterocytes, but largely dispensable in salivary gland cells for apical VSV G trafficking and basal GP64 trafficking. By examining host proteins potentially involved in protein trafficking in polarized cells, we identified clathrin adapter complexes AP-1 and AP-3, as well as seven Rab GTPases (Rab1, 4, 8, 10, 23, 30, and 35) that are important for directing basal trafficking of VSV G in polarized midgut enterocytes. We developed a new and powerful genetic system for these studies and answered fundamental questions regarding viral envelope protein trafficking in two critical tissues that serve as important barriers to systemic infection and transmission. However, our studies also raise many new questions regarding the mechanisms by which insect-specific viruses and arboviruses interact with and navigate through the tissues of their insect hosts.

## Acknowledgements

The authors thank Gary Whittaker (Cornell University) for providing anti-VSV G antibody 8G5F11, Mikio Furuse (NIPS) for antibodies directed against Snakeskin and Tetraspanin 2A, and Bruce Edgar for E-Cadherin-GFP *Drosophila* line. We also thank Peter Nagy for assistance and expertise with initial Rab GTPase localization mapping.

## Author Contributions

Conceptualization: Nicolas Buchon, Gary Blissard, *Jeffrey Hodgson*.

Formal analysis: Jeffrey Hodgson, Robin Chen, Nicolas Buchon, Gary Blissard

Funding acquisition: Gary Blissard, Nicolas Buchon.

Investigation: Jeffrey Hodgson, Robin Chen, *Nicolas Buchon, Gary Blissard*.

Methodology: Jeffrey Hodgson, Robin Chen, Nicolas Buchon, Gary Blissard.

Project administration: Gary Blissard, Nicolas Buchon.

Supervision: Nicolas Buchon, Gary Blissard.

Writing – original draft: Jeffrey Hodgson, *Robin Chen*.

Writing – review & editing: Jeffrey Hodgson, Robin Chen, Gary Blissard, Nicolas Buchon.

## Supplemental Data

**Figure S1.** The method for quantifying relative viral envelope protein distribution along the basal-apical axis in *Drosophila* midgut enterocytes and salivary gland cells. (A) Regions of interest (ROI, shown in yellow) from which signal intensity was measured along the basal-to-apical axis using ImageJ (as described in the Materials and Methods section). (B) Distribution of select marker proteins in midgut enterocytes and salivary gland cells. The peak in Discs large signal in enterocytes at the 52.5% mark (vertical dashed line) represents the location of septate junctions that define the boundaries between the basal and apical compartments. Phalloidin staining was used to assess the distribution of cortical actin in the salivary gland cells, indicative of the apical membrane invaginations. Phalloidin staining pattern and signal distribution closely approximate VSV G signal distribution (Fig. 2C and S3C), suggesting co-localization. The average signal distribution density is indicated by the horizontal dashed line and data points above the horizontal dashed line show regions of marker protein enrichment. (C) Locations of midgut enterocyte and salivary gland cell nuclei were determined by DAPI (blue line) signal distribution. The regions selected for statistical analysis to detect changes in viral envelope protein distribution are indicated by gray boxes. Those regions were selected based on average locations of marker proteins and nuclei (see the Materials and Methods section for details).

**Figure S2.** Levels of VSV G^WT^ and VSV G^ΔY^ signal in *Drosophila* midgut enterocytes at basal membrane surfaces and in the entire cell. (A) Non-permeabilized cells were used to determine the steady state levels of VSV G^ΔY^ (ΔY) compared to VSV G^WT^ (WT) at the basal membrane surface. Control cells that do not express any VSV G construct (N) were used to account for autofluorescence. Levels of VSV G^ΔY^ at the basal cell surface were substantially lower (11.4 times) than that of VSV G^WT^ (Pairwise Wilcoxon rank sum test: p = 2.6 × 10^-12^). B) Permeabilized cells were used to determine whether the ΔY substitutions affect steady state levels of VSV G^ΔY^ compared to VSV G^WT^ in the entire cell. Total levels of VSV G^ΔY^ in enterocytes were significantly lower (18.5 times) than that of VSV G^WT^ (Pairwise Wilcoxon rank sum test: p < 2.0 × 10^-16^). VSV G expression was induced for 3 days under the *da-Gal4 tub-Gal80^ts^* driver. VSV G was labelled with mouse anti-VSV G primary antibody and Alexa^647^-conjugated donkey anti-mouse secondary antibody. Signal levels were measured using ImageJ (as described in Materials and Methods). Since VSV G signal level differences could be the results of less efficient translation, processing, or turnover of the VSV G^ΔY^ protein, or from variations between different fly lines, we were unable to draw conclusions based on steady state levels of VSV G on the basal membrane surface or in the entire cell, but instead focused on the relative distribution patterns of the VSV G^WT^ and VSV G^ΔY^ proteins.

**Figure S3.** Cellular structural marker proteins in *Drosophila* midgut enterocytes and salivary gland epithelial cells. Tissues from flies constitutively expressing GFP-tagged E-Cadherin (labelling adherens junctions) were dissected, fixed and immunostained for Discs large (labelling septate junctions), Snakeskin (labelling septate junctions), Integrin β1 (labelling basal membrane), Tetraspannin 2A (labelling septate junctions), phalloidin (labeling cortical actin and visceral muscles), and DAPI (labeling DNA in nuclei). Scale bars represent 5 µm.

**Figure S4.** Localization patterns of VSV G in *Drosophila* salivary gland cells are variable. On the left, a bar graph illustrates the proportion of salivary gland cells displaying either basal, non-polarized, or apical enrichment patterns of VSV G^WT^ (n = 160) and VSV G^ΔY^ (n = 211). Basal enrichment pattern was found in 18.8% and 4.27% of salivary gland cells expressing VSV G^WT^ and VSV G^ΔY^, respectively. Non-polar distribution pattern was found in 52.5% and 54.5% of salivary gland cells expressing VSV G^WT^ and VSV G^ΔY^, respectively. Apical distribution pattern was found in 28.8% and 41.2% of salivary gland cells expressing VSV G^WT^ and VSV G^ΔY^, respectively. Microscopy images on the right illustrate examples of each pattern of VSV G localization. A vast majority of the salivary gland cells (81.3% for VSV G^WT^ and 95.7% for VSV G^ΔY^) showed presence of VSV G in non-basal regions. VSV G was labelled with mouse anti-VSV G primary antibody and Alexa Fluor 647 donkey anti-mouse secondary antibody and nuclei were stained with DAPI. Scale bars represent 5 µm.

**Figure S5.** Anthranilic acid inhibits basal VSV G trafficking in *Drosophila* midgut enterocytes. At 3-5 days post-eclosion, flies carrying an inducible VSV G expression construct (*Myo-Gal4, UAS-nlsGFP, tub-Gal80^ts^; UAS-VSV G^WT^*) were exposed to anthranilic acid-treated artificial diet (ACA; 240 mg dissolved in 300 μL of 100% ethanol applied onto the surface of ∼10 mL of artificial diet and allowed the ethanol vehicle to completely evaporate before introducing the flies) or the ethanol vehicle-treated diet (ETH; 300 μL of 100% ethanol) for 3 days at 18°C before shifting to 29°C for 48 h to induce VSV G expression. The average relative position of the nucleus was also plotted as a gray line. The average cell height (basal-to-apical distance in %) (x-axis) and signal intensities (y-axis) were measured and calculated as described in the Materials and Methods section. The basal-to-apical distribution of VSV G resulting from each treatment is shown on the graph. The basal and apical regions on the graphs were established based on the location of the septate junction (discs-large polarity marker) (Fig. S1), and the division is indicated at 52.5% on the x-axis by a vertical dashed line. The shaded areas on each graph represent the areas of the cell (basal 20% and apical 40%) relatively free from influence of the nuclei and used for statistical analyses. The number of asterisks refer to the level of statistical significance (n.s. p > 0.05, * p ≤ 0.05, ** p ≤ 0.01, *** p ≤ 0.001, and **** p ≤ 0.0001) between distributions of VSV G in the midgut enterocytes of ACA (n = 68) and ETH (n = 80) treated flies in the shaded areas. Data shown represents combined data from 3 independent replicates.

**Figure S6.** Distribution patterns of YFP-tagged Rab GTPases in *Drosophila* midgut enterocytes and salivary gland cells. Fly lines (Zhang et. al 2007. Genetics 176:1307-22) inducibly and ubiquitously expressing each of 30 UAS-YFP-Rab GTPases for 3 days using the *da-Gal4 tub-Gal80^ts^*driver were immunostained with chicken anti-GFP primary antibody and Alexa Fluor 488 goat anti-chicken secondary antibody (green). Actin (red) was labelled with Alexa Fluor 555 phalloidin, nuclei (blue) were stained with DAPI, and Discs large (magenta) was stained with mouse anti-Discs large primary antibody and Alexa Fluor 647 donkey anti-mouse secondary antibody. Scale bars represent 5 µm.

**Table S1.** Recipe of artificial *Drosophila* diet used in this study.

